# Electrophysiological analysis of healthy and dystrophic 3D bioengineered skeletal muscle tissues

**DOI:** 10.1101/2020.11.10.376764

**Authors:** Christine T Nguyen, Majid Ebrahmi, Penney M Gilbert, Bryan A Stewart

## Abstract

Recently, methods for creating three-dimensional (3D) human skeletal muscle tissues from myogenic cell lines have been reported. Bioengineered muscle tissues are contractile and respond to electrical and chemical stimulation. In this study we provide an electrophysiological analysis of healthy and dystrophic 3D bioengineered skeletal muscle tissues. We focus on Duchenne muscular dystrophy (DMD), a fatal muscle disorder involving the skeletal muscle system. The *dystrophin* gene, which when mutated causes DMD, encodes for the Dystrophin protein, which anchors the cytoskeletal network inside of a muscle cell to the extracellular matrix outside the cell. Here, we enlist a 3D *in vitro* model of DMD muscle tissue, to evaluate an understudied aspect of DMD, muscle cell electrical properties uncoupled from presynaptic neural inputs. Our data shows that electrophysiological aspects of DMD are replicated in the 3D bioengineered skeletal muscle tissue model. Furthermore, we test a block co-polymer, poloxamer 188, and demonstrate capacity for improving the membrane potential in DMD muscle.Therefore, this study serves as the baseline for a new *in vitro* method to examine potential therapies directed at muscular disorders.

## Introduction

Muscular dystrophies (MDs) disorders of skeletal muscle that can be classified based on age of onset, overall progression, and mode of inheritance. MDs are characterized by progressive muscle weakness and muscle wasting. Histological analysis shows distinguishing features of MDs that include non-uniform fiber sizes and variations, areas showing muscle necrosis, and increased aggregates of connective tissue and fats (Reviewed by Emery, 1998).

Duchenne muscular dystrophy (DMD) is a fatal X-linked disorder caused by mutations of the *dystrophin* gene at the Xp21 locus (Koenig et al., 1987). Duchenne muscular dystrophy (DMD) is the most severe and most common form of muscular dystrophy. Patients with DMD demonstrate progressive muscle weakness and eventually die in the third decade of life (Straub and Campbell, 1997, Blake et al., 2002). DMD muscle fibers undergo episodes of regeneration and necrosis, and while regeneration is active, it is not enough to overcome necrosis as reformed muscle cells still contain the dystrophin mutation (Carpenter and Karpati, 1979).

Dystrophin is a 427-kDa cytoplasmic protein spanning ∼2.5Mb of the genomic sequence (Koenig, 1987). Studies have shown that Dystrophin is an important cytoplasmic protein that underlies the plasma membrane in skeletal muscle where it connects actin with a Dystrophin-associated glycoprotein complex (DGC) involved with overall muscle stability (Huard et al., 1992; Blake et al, 2002). In the absence of Dystrophin, as occurs in DMD muscle fibers, the plasma membrane becomes fragile, reducing the stiffness of the muscle, and increasing membrane leakiness and leading to exposure of the Dystrophin-deficient muscle fiber to hypoosmotic conditions and permeability (Goldstein and McNally, 2010).

In the last 30 years, there have been many different animal models of DMD ranging from invertebrates, such as *Drosophila melanogaster*, to the most widely used mammalian model, the *mdx* mouse (Reviewed by McGreevy et al., 2015). A cure for DMD could entail genome editing in which the *dystrophin* gene is corrected, using one of a number of putative strategies, such as exon skipping, or adeno-associated virus (AAV) based gene therapy enlisting the CRISPR/Cas9 system (Bengtsson et al., 2017; Shimizu-Motohashi et al., 2019; Moretti et al, 2020; Amoassi et al., 2018, Min et al., 2019), or otherwise (Mollanoori et al., 2020).

Since no strategy to date has been successful in correcting dystrophin mutations in the stem cell compartment, we expect that complimentary therapies will be needed. For example, membrane properties are being explored in pre-clinical studies as a putative DMD therapy approach, for example examination of lipid metabolism and its influence on plasma membrane properties on human cells in culture (Le Borgne, 2012). Given that DMD muscle membranes are fragile and susceptible to damage and disruption, investigations aimed at studying the electrical properties of the muscle membrane, and how using therapeutic reagents such as block co-polymers can change its electrical properties, are needed.

Increasingly, directed design of *in vitro* systems is playing a role to aid our understanding biological systems. Among the three-dimensional (3D) platforms for *in vitro* studies, a functional model for human skeletal muscle tissue established from the differentiation of primary human myoblast cell lines (Madden et al., 2015) and immortalized myoblast cell lines (Mamchaoui et al., 2011) have been described

3D culture models are thought to more accurately represent native muscle because the cellular and physical (Lutolf et al. 2009; Morrissey et al. 2015) microenvironments are known to exert regulatory control over skeletal muscle tissue homeostasis (Gilbert et al. 2010; Quarta et al. 2016), regeneration (Cosgrove et al. 2014), and disease pathogenesis (Urciuolo et al. 2013). In addition to an increased lifespan *in vitro*, 3D muscles have been shown to have larger acetylcholine receptor (AChR) clusters, as well as an increase in the expression of adult isoforms of myosin heavy chain proteins as compared to 2D muscle cultures (Bakooshli et al., 2019). One goal of 3D DMD muscle model studies is to find therapeutic targets for the disease through drug screens and pathogenetic studies. Currently, 3D platform models of skeletal muscles using iPSCs have been developed for disease modeling, including DMD (Maffioletti et al., 2018; Piga et al., 2019). However, the electrophysiological properties of the muscle cells within 3D skeletal muscle tissues established in culture have yet to be characterized.

In this study we report the electrophysiological characteristics of myotubes within 3D bioengineered skeletal muscle cells derived from immortalized myoblast cell lines using sharp microelectrode techniques. We generated 3D bioengineered skeletal muscle tissues using the immortalized DMD cell lines DMD5-13515, and KM571-DMD (Mamchaoui et al., 2011). The resting membrane potential (RMP), the membrane resistance, and muscle excitability were measured and compared to myotubes within 3D muscle tissues derived from a healthy human immortalized myogenic cell line. We show that DMD skeletal muscle cells are more depolarized, and that membrane integrity, measured by membrane resistance, is compromised in DMD cells when compared to healthy muscle cells. We further demonstrate that the RMP of dystrophic myotubes becomes more negative upon incubating DMD muscles with poloxamer 188, suggesting the treatment improves the physiological condition of the muscle cell.

## Materials and Methods

### Immortalized human myoblast propagation

The AB1167, DMD-5-13515, and KM571 immortalized human myoblast cell lines were received from Vincent Mouly and Anne Bigot (Institut de Myologie, Paris, France) who established the lines as previously described (Mamchaoui et al., 2011). AB1167 was derived from fascia lata muscle of a healthy 20-year old male, DMD-5-13515 from paravertebral muscle of a 14-year old male with DMD a dystrophin gene duplication in exon 2, and KM571-DMD from fascia lata muscle of a 10-year old male with a dystrophin gene deletion of exon 52 (Mamchaoui et al., 2011). The immortalized myoblast cell lines were maintained on 10 cm tissue culture dishes containing 7 mL of skeletal muscle cell growth medium (79 % PromoCell media (C-23160) supplemented with 15 % fetal bovine serum (Gibco) and with 1 % penicillin-streptomycin (Life Technologies). Myoblast cells were kept at 37°C with 5 % CO_2_. Half of the growth media was exchanged every other day. Cells were passaged to a new plate upon reaching 90 % confluency.

### Generation of 3D bioengineered skeletal muscle tissues

Custom made 12-well culture plates containing a dumbbell shaped channel, to which Velcro™ was glued at each end, were prepared ahead of experiments exactly as described in (Bakooshli et al., 2019). Devices were sterilized with 70 % ethanol prior to use. Cell suspensions from cultures at passages under 10 (P<10) were used for the generation of three-dimensional (3D) human skeletal muscle tissues. 3D muscle tissues were prepared as previously described (Madden et al., 2015, and Bakooshli et al., 2019), with the addition that individual wells in the 12-well plate were incubated with pluronic acid (Sigma; P2443) overnight prior to tissue seeding. Briefly, cell suspensions at a density of 1,500,000 cells/tissue was mixed with a hydrogel mixture containing 4 mg/ml fibrinogen (40 % v/v; Sigma; F8630), 20 % Geltrex (Thermo Fisher Scientific), and 40 % Dulbecco’s modified eagle medium (DMEM; Thermo Fisher). Thrombin (Sigma; T6884) was added to the hydrogel/cell mixture at 0.2 unit per mg of fibrinogen just prior to seeding the hydrogel/cell mix. The hydrogel/cell mix containing thrombin was then evenly spread along the length of the channel. Tapping the edges of the 12-well plate ensured even dispersion of the mixture within the channel. The 12-well plate was then incubated at 37 °C for 5 minutes to allow for fibrin polymerization (Bakooshli et al., 2019). Skeletal muscle cell growth media containing 1.5 mg/mL 6-aminocaproic acid (ACA; Sigma) was gently added to each channel and then to fill the well. Two days later the skeletal muscle cell growth media was then substituted with myoblast differentiation media (97 % DMEM (Thermo Fisher), 2 % horse serum (Gibco), 1 % penicillin-streptomycin (Life Technologies), and 10 μg/mL insulin (Gibco) supplemented with 2 mg/mL 6-aminocaproic acid (ACA; Sigma). Tissues were maintained at 37 °C with 5 % CO_2_. Half of the differentiation media was exchanged every other day. In all experiments the 3D tissues established from AB1167 were differentiated for 8 or 10 days, whereas 3D tissues established from KM571-DMD, and DMD-5-13515 were differentiated for 8 days.

### Electrophysiology

On the appropriate day, differentiation media was aspirated from muscle tissues and replaced with HEPES buffered Tyrode’s solution [119 mM NaCl, 5 mM KCl, 25 mM HEPES buffer, 2 mM CaCl_2_, 2 mM MgCl_2_, 6 g/liter glucose (Cold Spring Harbor Protocols)] at room temperature. The RMP and muscle membrane resistance was measured in current clamp, using a sharp intracellular microelectrode backfilled with 3M KCl. Glass intracellular microelectrodes were pulled using a micropipette puller (Sutter Instruments), and had a resistance of ∼30 MΩ. Electrophysiological measurements were obtained with an AxoClamp 2B amplifier (Axon Instruments) to amplify analog signals, digitized Digidata 1322A (Axon Instruments) and recorded with pClamp8.2 software. The recordings were analyzed using pClamp 8.2 as well as Clampfit 10.0 software.

Depolarization of muscle cells to study excitation was achieved using two methods: direct current injection and by anode break excitation. The amplitude of active responses was measured using the difference of two cursors, one paced at the baseline of the event and one at the peak of the event in Clampfit 10.0 software. The cursors were locked at a distance from each other by a standardized value of 9.0 ms. All data exported to GraphPad Prism 5 for statistical and graphical analysis.

To examine the ionic contributions to excitation, NaCl was substituted with equimolar choline chloride in one experiment, and Ca^2+^free solution with the addition of EDTA was used for another.

### Poloxamer treatment

3D skeletal muscle tissues were incubated for 1 hour in poloxamer 188 (P188; Sigma, P5556) at 37 °C. Poloxamer was diluted to a final concentration of 1 % in differentiation media. Following incubation, poloxamer and differentiation media was aspirated and washed with HEPES buffered Tyrode’s solution. For recordings, HEPES buffered Tyrode’s solution was used to fill the well with the muscle tissue.

### Immunohistochemistry

3D bioengineered skeletal muscle tissues were fixed for 20 minutes in 4 % paraformaldehyde (Ted Pella, Inc) in phosphate buffered saline (PBS). Muscle tissues were then washed with PBS containing 0.1 % Triton X-100 (PBT). Muscle tissues were then placed into a microcentrifuge tube containing PBT with 2 % final concentration of normal goat serum (NGS; Vector Labs) for 30 mins. Blocking solution was removed and replaced with PBT.

Monoclonal Anti-α-Actinin (Sarcomeric) antibody (Sigma, A7811 clone EA-53) at a dilution of 1:100 was added to the PBT and allowed to incubate overnight at 4 °C with rocking. Following incubation with primary antibody, muscle tissue was washed with PBT at room temperature 3 times for 20 minutes. Muscle tissue was then mounted on a glass slide in Vectashield mounting medium for fluorescence with DAPI (Vector Labs). A cover slip was placed over the muscle tissue. All images were collected on a Zeiss LSM 800 confocal microscope; images were taken using a 20x/0.8 air or a 40x/1.4 oil immersion lens.

### Transmission Electron Microscopy

Muscle tissues were fixed in 2.5 % glutaraldehyde and 2 % paraformaldehyde in 0.1 M sodium cacodylate buffer for 2 hours at room temperature. Sample embedding and sectioning was completed at the Nanoscale Biomedical Imaging Facility at The Hospital for Sick Children, Toronto Canada. Muscle tissue samples were washed in 0.1 M sodium cacodylate buffer with 0.2 M sucrose, and fixed for 1.5 hours on ice in 1 % osmium tetroxide and washed in cacodylate buffer. An alcohol series of dehydration occurred where muscle samples were washed three times in 100 % ethanol, and then washed twice in propylene oxide. Muscle samples were then placed in 50:50 propylene oxide/Quetol-Spurr resin, then in 100 % resin for two hours. Muscle samples were then embedded in 100 % resin overnight at 65 °C. Muscle samples were then sectioned longitudinally and with cross-sections. Muscle samples were collected on mesh grids and stained with lead citrate and uranyl acetate. Images were processed and collected on a FEI Technai 20 electron microscope.

### Statistics

In this study, immortalized myoblast cell lines from one healthy (AB1167) and two dystrophic (KM571-DMD and DMD5-13515) muscle patient donors were used. For the DMD tissues, four independent experiments from each muscle cell line were performed, where each experiment included 3 muscle tissues, for a total of 12 muscle tissues per muscle line (*n* = 12 tissue replicates). For the healthy tissue, 2 independent experiments were performed with a total of 6 muscle tissues assessed (*n*=6 tissue replicates). Statistics were performed on the mean values of individual myotubes impaled and recorded on from n=12 DMD muscle tissues and n=86 healthy muscle tissues. For experiments investigating poloxamer 188 treatment, a single independent experiment was performed using the AB1167 line (day 10), and the DMD5-13515 line (day 8), where a total of 3 tissues for each muscle line was assessed (*n*=3 tissue replicates). For experiments studying the effect of ion composition, one independent experiment was performed on the AB1167 line (day 10), with a total of 3 tissues (*n*=3 tissue replicates). Statistical analysis was performed using GraphPad Prism 5.0 software. Unpaired t-test as well as one-way ANOVA with Tukey and Bonferroni post-test was used for statistical differences and comparisons

## Results

### Resting membrane potential of healthy and DMD muscle in 3D culture

We first conducted studies to compare the RMP of muscle cells established using immortalized myoblasts from healthy individuals to those harbouring mutations in the *dystrophin* gene. The mean RMP for healthy myotubes on day 10 of differentiation was determined to be −74.3 ± 0.6 mV (*n=*41 different muscle cells from 12 different muscle tissues).

Next, 3D muscle tissues derived from DMD cell lines KM571-DMD, and DMD5-13515 were evaluated. We found that the RMP of both cell lines was significantly more positive than that of the healthy tissue. Specifically, KM571-DMD myotubes had a RMP of −41.5 ± 2.4 mV (*n=*27), and DMD5-13515 myotubes had a RMP of −44.2 ± 2.3 mV (*n=*22) (Fig. 1).

**Figure 1.**
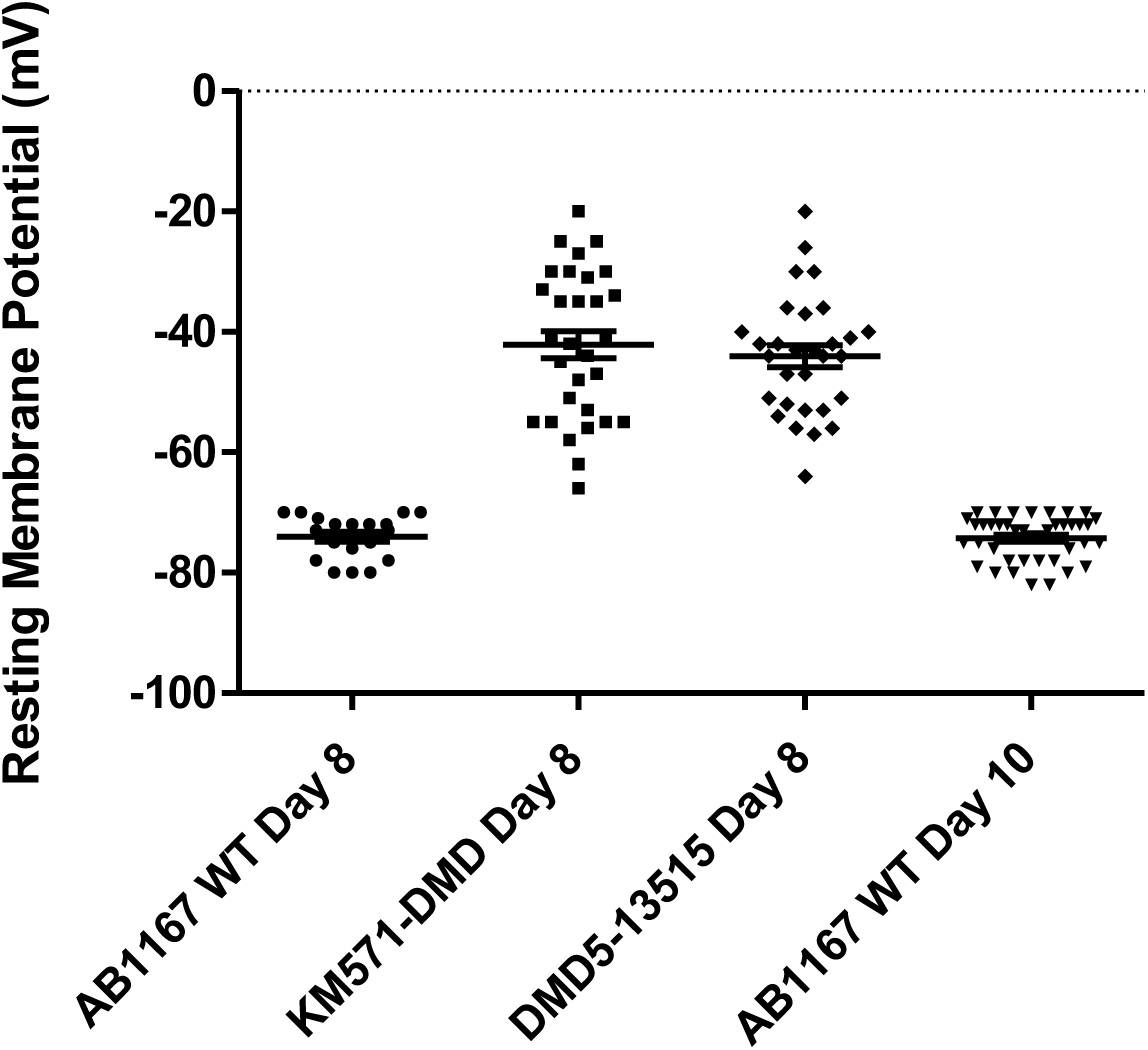
The resting membrane potential of DMD myotubes are more positive than healthy myotubes. Each point in the figure represents a single myotube measurements from 12 different muscle tissue for each muscle line, except for AB1167 day 8 where data is from 6 muscle tissues (*n*=individual myotubes). The mean ± s.e.m RMP of muscle cells are shown by the lines and error bars. KM571-DMD and DMD5-13515 day 8 RMPs are significantly more positive than both AB1167 day 8 and AB1167 day 10 (one-way ANOVA Bonferroni’s post-test, *p*<0.0001).

As the integrity of DMD muscle tissues was not sustained to day 10 of differentiation, (e.g. myotubes frequently broke by this time-point), the DMD cell lines were recorded on day 8 of differentiation. Therefore, we also analyzed myotubes in AB1167 muscles on day 8 of differentiation, and found that the mean RMP at this time-point was similar to the day 10 values, and which was significantly more negative than DMD cell lines. The mean RMP for AB1167 WT at day 8 post differentiation was −66.4 ± 6.7 mV (*n=*20). These findings suggest that the DMD muscle cells do not possess a healthy physiological RMP as would be expected for proper muscle signaling and function.

### Membrane resistance is disrupted in DMD myotubes

We next evaluated the membrane resistance of healthy and dystrophic myotube. Current was injected in increments of 0.2 nA. Based on 1 nA of current injection, healthy myotubes in 3D muscles on day 10 of differentiation exhibited a mean muscle membrane resistance of 66.5 ± 6.7 MΩ (*n=*9; Fig.2) on day 8 and 74.0 ± 8.8 MΩ (*n=*8; Fig.2) on day 10.

**Figure 2.**
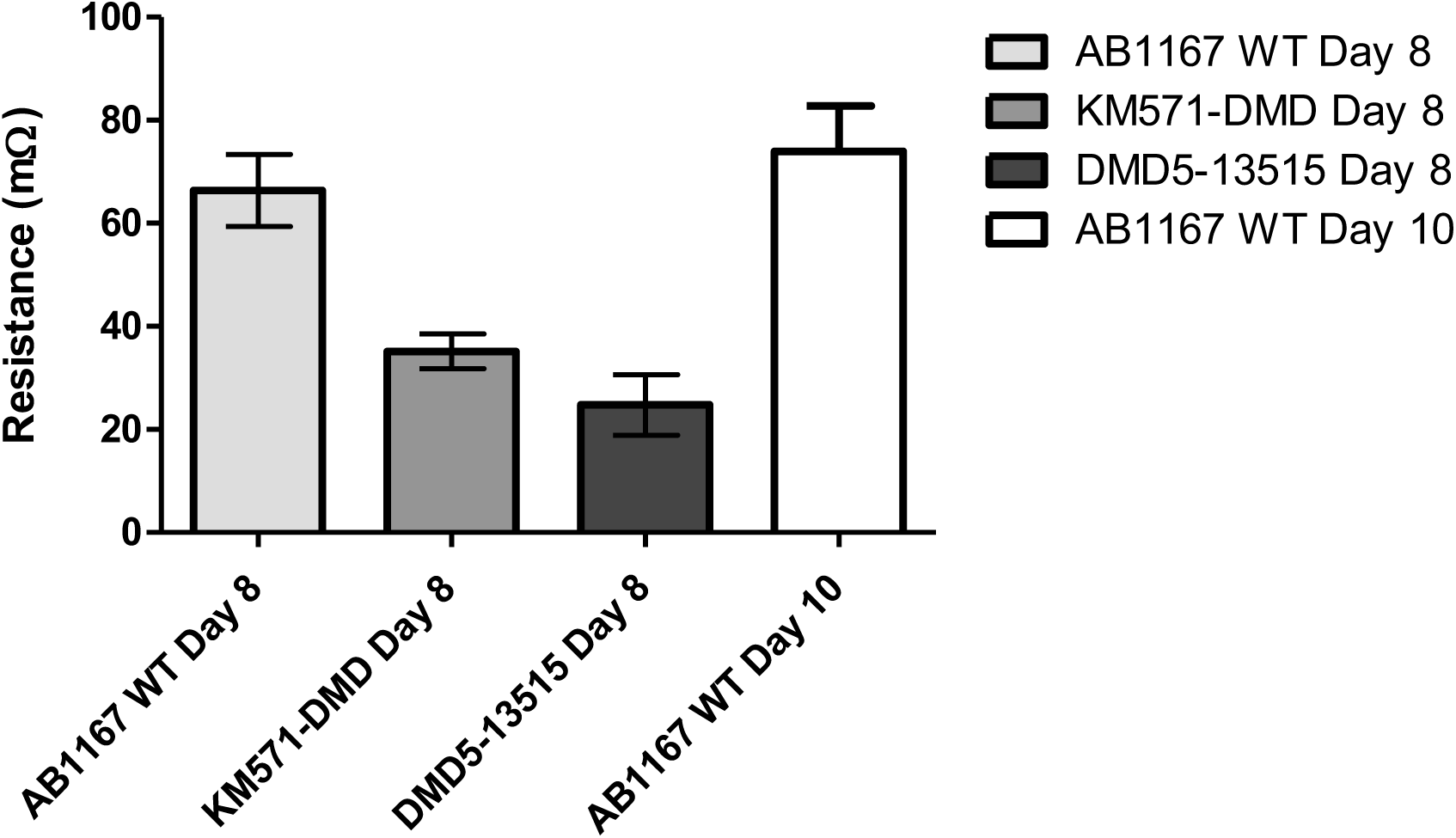
Healthy myotubes have a higher muscle membrane resistance when compared to DMD myotubes. The muscle membrane resistance was measured on individual myotubes within 3D bioengineered skeletal muscle tissues. The bars represent the mean ± s.e.m membrane resistance from single myotube measurements of 12 different muscle tissue for each muscle line, except for AB1167 day 8 where data is from 6 muscle tissues (*n*=individual myotubes recordings). KM571-DMD \*\**p*<0.01, DMD5-13515 \*\*\*\**p*<0.0001, one-way ANOVA Bonferroni’s post-test (day-8). There is no significant difference between the membrane resistance of KM571-DMD and DMD5-13515.

We next evaluated the muscle membrane resistance of DMD tissue and we found that the membrane resistance of DMD muscle tissues day 8 post differentiation was significantly lower that of healthy muscle tissue. Specifically, KM571-DMD myotubes had a mean membrane resistance of 35.2 ± 3.4 MΩ (*n=*10; Fig. 2), and DMD5-13515 myotubes had a mean membrane resistance of 24.8 ± 5.9 MΩ (*n=*13; Fig. 2). There was no significant difference between the membrane resistance of KM571-DMD and DMD5-13515. These findings suggest that there are disruptions of the plasma membrane of DMD myotubes that affect the electrical activity of the cell.

### Dystrophic myotubes within 3D bioengineered skeletal muscle tissues lack excitability

In order to evaluate the excitability of myotubes in healthy muscle as compared to dystrophic muscle, we sought to electrically stimulate muscle cells with injected current using a sharp microelectrode and study active responses. The term “active response” is used because the recorded muscle excitability responses do not represent the “all or none” representation of an action potential; however, there is still a rise and falling phase. Active responses, or muscle excitability, was achieved from myotubes within 3D bioengineered skeletal muscle tissue using two methods. The first method used a bridge circuit that allowed for a single microelectrode to inject current, also while recording simultaneously. Representative active response traces using bridge circuit can be seen in Figure 3. The second method that was used to achieve muscle excitability through active responses was by using anode break excitation (Katz, 1939; Guttman and Hachmeister, 1972). Small action potentials generated using the anode break method have been observed in rat skeletal muscles (Kidokoro, 1973). Representative active response traces using anode break excitation of myotubes can be seen in Figure 3. When comparing muscle excitability in healthy myotubes and DMD myotubes, excitability in the presence of active responses was frequent in healthy muscle, at both days 8 and 10 of differentiation. The presence of active responses from healthy muscle cells had variable amplitudes measured from anode break traces, however could not be compared with DMD muscle cells as there was no active responses achieved using anode break excitation with DMD muscle cells.

**Figure 3.**
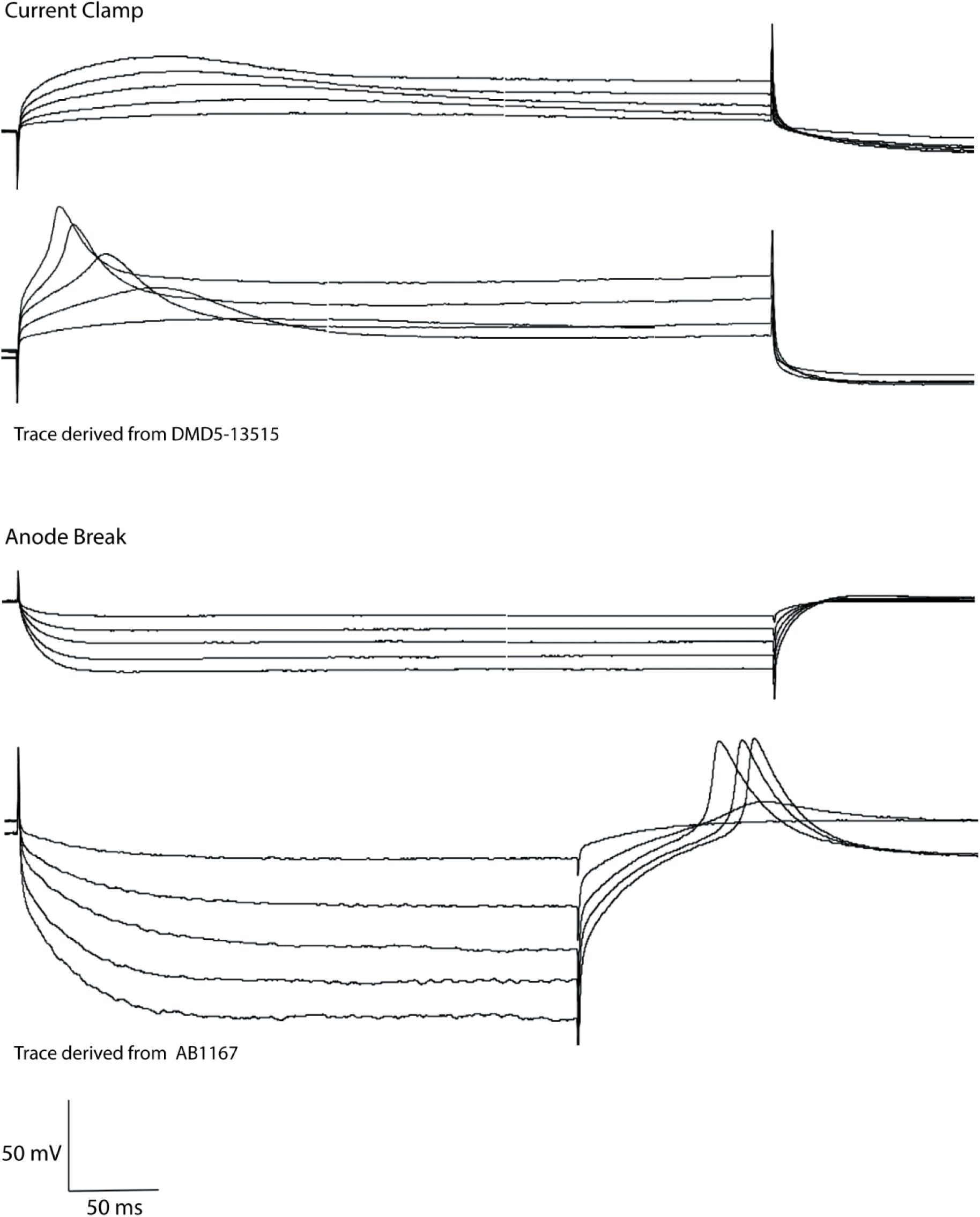
Representative traces of muscle excitability in 3D bioengineered human skeletal muscle. Active responses are shown using two electrical stimulation methods: current clamp (top), and anode break (bottom). Electrical stimulation is shown with increasing step-wise increments of 0.2 nA of injected current, from 0.2 nA to 1 nA of current injection.

### Active responses from muscle cells within healthy tissues are Na^+^ dependent

Since this the first report of electrical activity of 3D cultured muscle, we sought to understand the ionic basis of the excitable response. Accordingly, Na^+^ was replaced with equimolar choline chloride in one experiment and Ca^2+^ was replaced with EDTA in another. Replacing Na^+^ with choline chloride in either calcium containing (*n*=5), and calcium free solutions *(n*=6) resulted in the absence of any active responses. No depolarizations were seen (Fig. 4). Removing calcium and adding EDTA in the buffer solution had no effect on muscle excitability, as active responses were still present. These results suggest that active responses from the healthy muscle cells are Na^+^ induced, and they are not affected by Ca^2+^. The presence of Na^+^ is needed for muscle cell excitability.

**Figure. 4.**
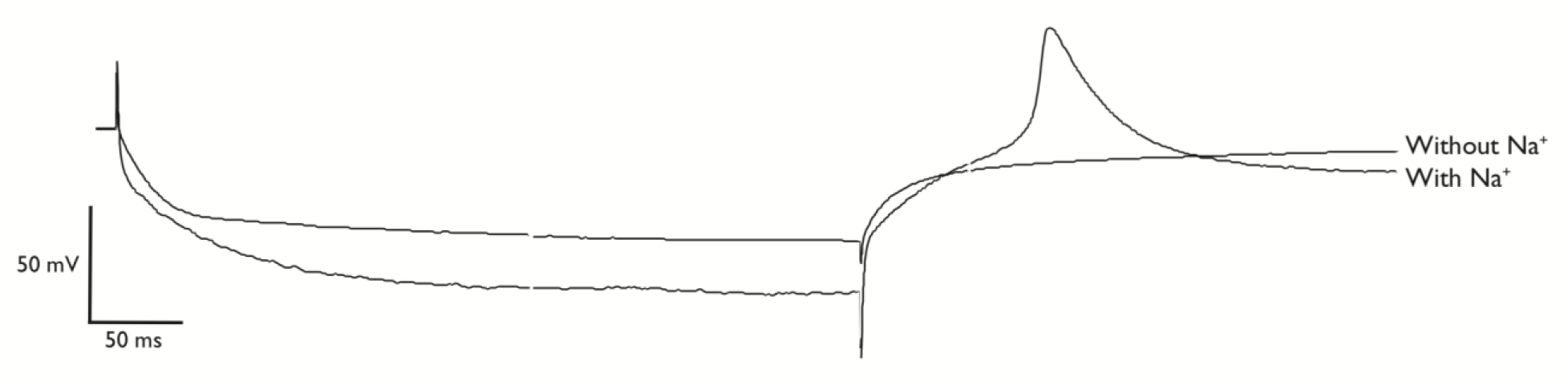
Representative trace of active response in the presence or absence of external Na^+^. In the absence of Na^+^, there is no active response present. Traces are shown with 0.6 nA of injected current. Trace shown was collected from muscle cells derived from immortalized cell line AB1167. In all six tissues with choline chloride (*n*=6), no active response was seen. In all four (*n*=4) muscle tissues without choline chloride, active traces were seen.

**Figure 5.**
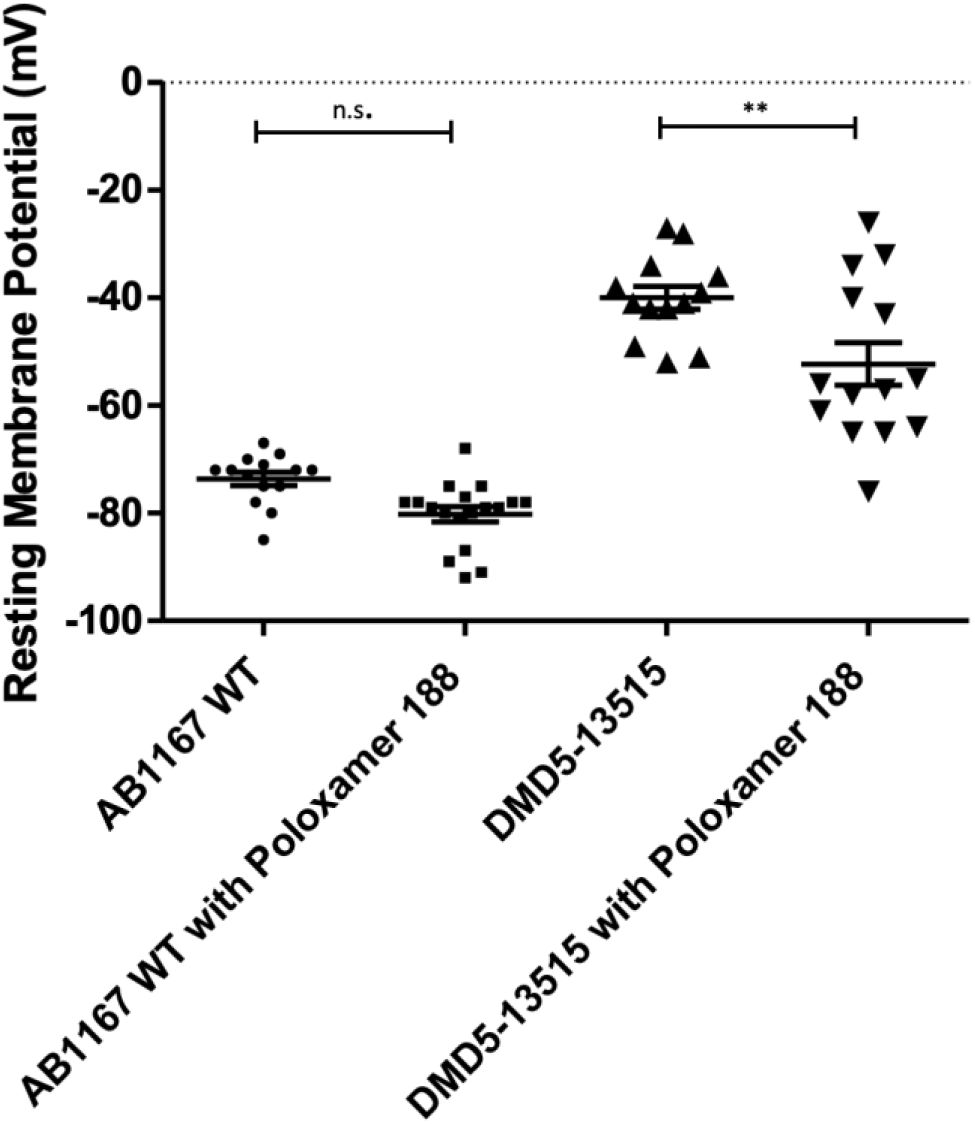
Incubation with poloxamer 188 leads to hyperpolarization of the resting membrane potential of dystrophic myotubes within 3D bioengineered skeletal muscle. Muscle tissues were incubated with 1 % poloxamer188 for 1 hour at 37 °C. The RMP was measured where each point represents the RMP for a single myotube obtained from 3 muscle tissues for both AB1167 and DMD5-13515 (one-way ANOVA Tukey post-test, not significant (*n*.*s*.); \*\**p*<0.001). AB1167 WT (*n*=14), AB1167 with P188 (*n*=18), DMD5-13515 (*n*=13), and DMD5-13515 with P188 (*n*=14). Muscle tissues used were from day 10 post differentiation for AB1167, and day 8 for DMD5-13515.

### Poloxamer 188 hyperpolarizes the resting membrane potential of dystrophic myotubes

Since muscle membrane leakiness is an important component of the DMD phenotype, we next used a biomaterial approach to determine if we could improve the membrane resistance of DMD muscle cells. Specifically, muscle tissues were incubated with 1 % of poloxamer 188 (P188) for 1 hour at 37 °C. The RMP was determined for the healthy tissue and the dystrophic DMD5-13515 cell line. Incubation with the P188 led to a significantly more negative RMP of the DMD muscle cells (\*\**p*<0.001; one-way ANOVA, post Tukey test). The mean RMP of DMD muscle cells without P188 was −45.50 mV ± 2.2 mV. Following incubation with P188 the RMP was more negative, with a mean of −64.25 mV ± 3.9 mV. This brought the RMP of DMD muscle cells much closer to the healthy physiological RMP and suggests that P188 could improve membrane disruptions present in DMD muscle cells, thereby allowing it to come to a RMP closer to that of healthy myotubes.

### Myotube Size

Next, we conducted morphological analyses with which to relate our electrophysiological analyses. Using immunohistochemistry, 3D skeletal muscle tissues were stained to visualize sarcomeric α-actinin and with this the width of individual myotubes was determined. Here, we show that the myotubes within DMD muscles tissues have a smaller width than those in healthy muscle tissues (Fig. 6). The mean width of myotubes in healthy tissue is 10.5 ± 0.3 μm (*n=*30), 8.3 ± 0.3 μm (*n=*30) for KM571-DMD tissue, and 9.8 ± 0.6 μm (*n=*30). KM571-DMD day 8 is significantly different from AB1167 day 10 (\*\**p*<0.001) and DMD5-13515 day 8 *(*p*<0.1). AB1167 day 10 and DMD5-13515 have no significant differences. Together, this suggests that aberrant electrical properties are not always associated with diminished in myotube width.

**Figure 6.**
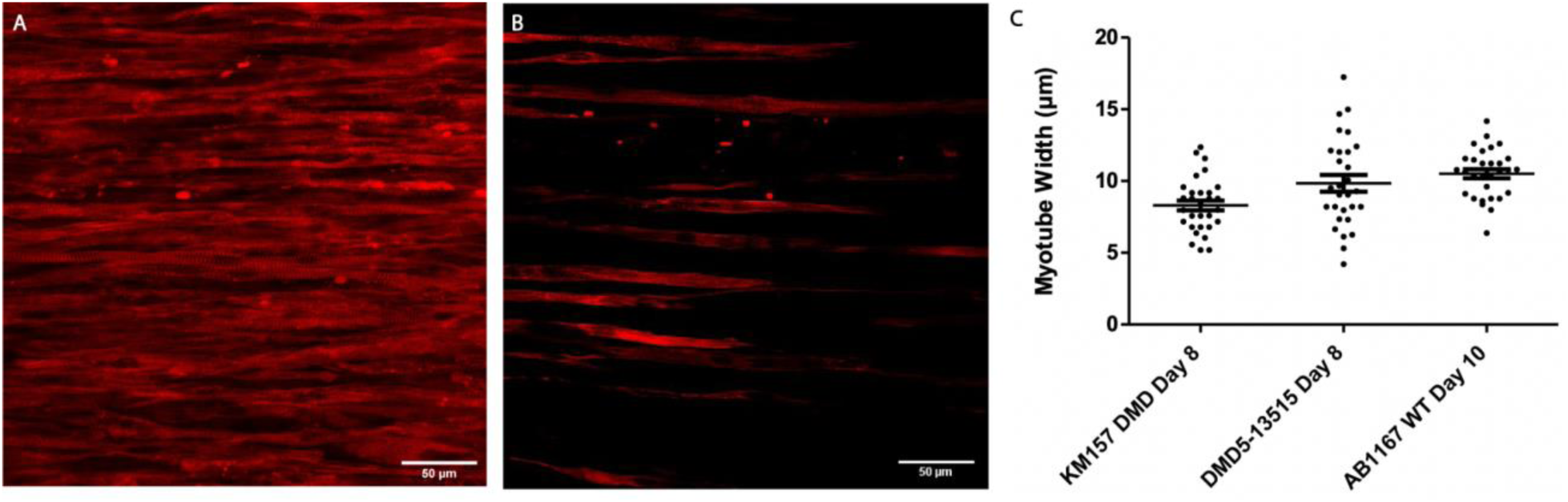
Myotubes of healthy tissues are wider than DMD myotubes. Using immunohistochemistry, myotubes were stained to visualize sarcomeric α-actinin (Abcam, ab9465). A) Healthy muscle tissue from AB1167 day 10-post differentiation. Scale, 50μm. B) Dystrophic muscle tissue KM571-DMD day 8-post differentiation. Scale, 50μm. C) Myotubes widths were determined using the line tool on ImageJ. Each point on the figure represents a single myotube (*n*=30 myotubes for each muscle line). KM571-DMD day 8 is significantly different from AB1167 day 10 (\*\**p*<0.001) and DMD5-13515 day 8 *(*p*<0.1) (One-way ANOVA; Bonferroni post-test).

### DMD muscle cells are disorganized at the ultrastructural level

Next, we examined myotube ultrastructure to determine whether there are differences between healthy and DMD myotubes that may explain the aberrant membrane properties we measured of DMD myotubes. Transmission electron microscopy (TEM) was used to compare between healthy and DMD 3D bioengineered skeletal muscle tissues. Sarcomere organization was visible in the AB1167 WT muscle cells, such that muscle bands discs were clearly identifiable (Fig. 7). On the other hand, myotubes within DMD muscle tissues showed sarcomere disorganization. Repeating units of the sarcomere consisting of Z-disk and bands were not observed in dystrophic muscle (Fig. 8). The apparent disorganization of fibers seen in DMD muscles are indicative of unhealthy muscle physiology. The membrane properties measured from DMD myotubes could be attributed to the lack of skeletal muscle organization seen in the diseased muscle, which could be the cause for the non-physiological recorded measurements in DMD electrophysiological recordings.

**Figure 7.**
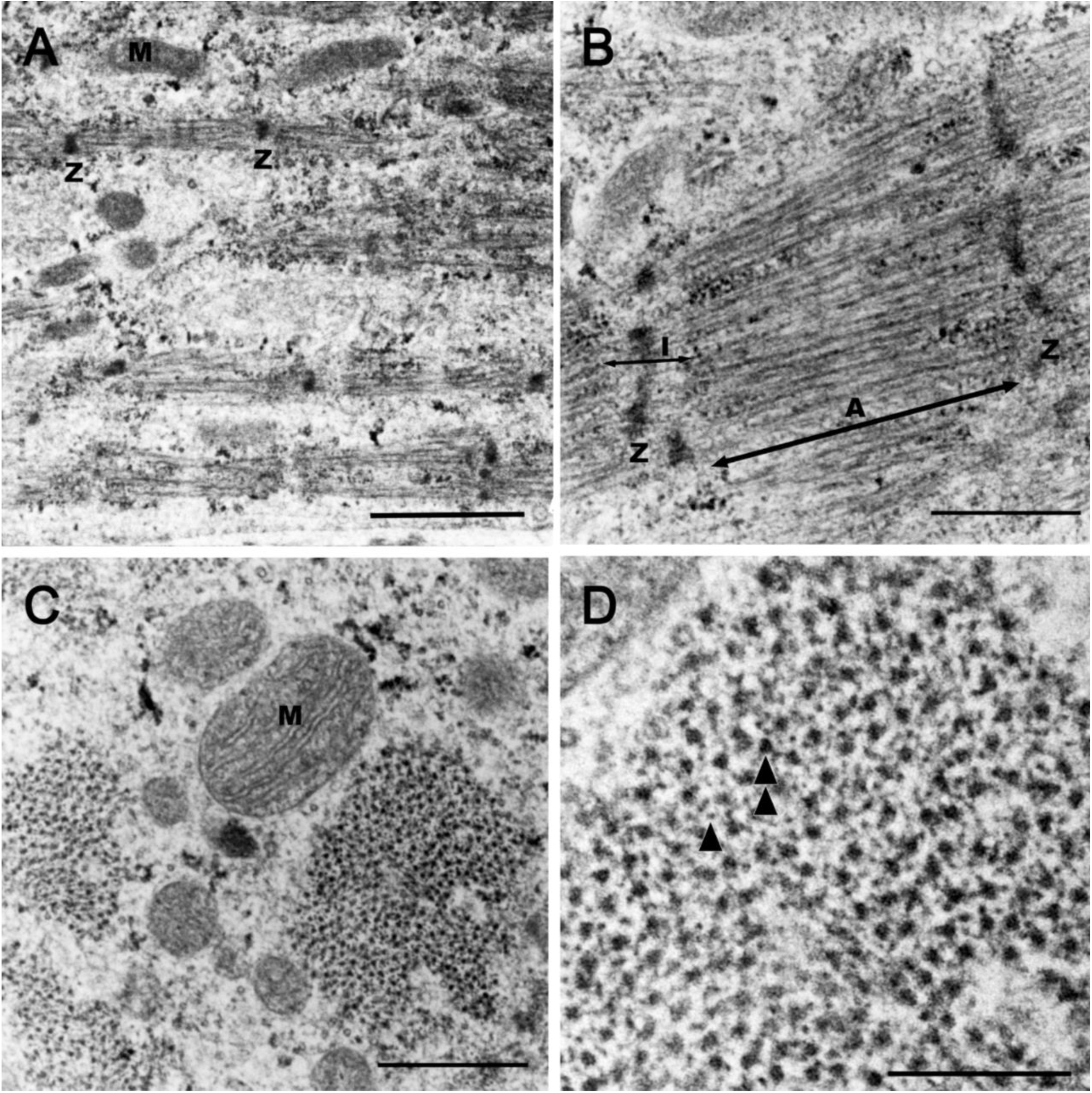
Ultrastructure of 3D bioengineered human skeletal muscle from human immortalized cell line AB1167 WT on 10 day of differentiation. (A) Longitudinal section showing fiber organization with sarcomeric z-line-Z, and mitochondria-M [Scale bar, 1 µm]. (B) Longitudinal section showing sarcomere with A band, I band, and Z line [Scale bar, 500 nm]. (C) Cross section showing myofibrils and filaments with mitochondria-M [Scale bar, 500 nm]. (D) Cross section showing the different filaments where thin filaments are shown with a single arrowhead, and thick filaments is shown with a double arrowhead [Scale bar, 200 nm].

**Figure 8.**
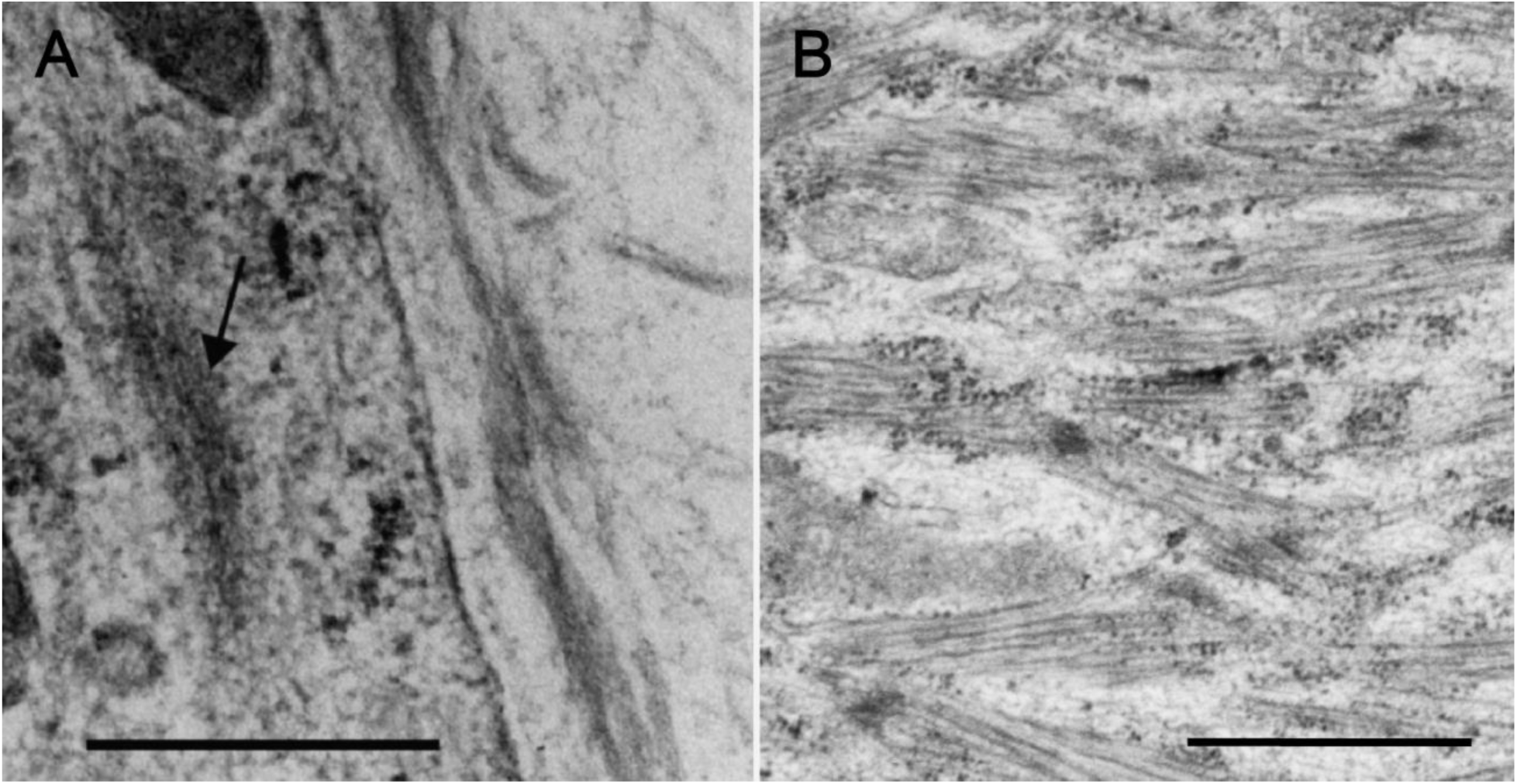
Ultrastructure of 3D bioengineered human skeletal muscle from immortalized DMD cell lines 10-day post-differentiation. (A) Disorganized fibers seen with arrow from DMD cell line KM571-DMD. Scale, 500 nm. (B) Disorganized sarcomeres without banding or Z-disc visible. Muscle fibers shown from DMD cell line DMD5-13515. Scale, 1µm.

## Discussion

This study provides the first physiological analysis of bioengineered muscle tissue grown in a 3D culture system. The aims of the study were threefold: to establish baseline parameters for healthy muscle; to compare healthy muscle to DMD muscle; and to provide proof of concept for the therapeutic screening potential of this system, for DMD and other muscle disorders. The key findings of our work are that the data we obtained for healthy muscle largely agree with that of the published literature, several key DMD phenotypes are recapitulated in the 3D cultures, and incubation of DMD tissue in poloxamer 188 can partially restore DMD phenotypes towards normal. Together with our previous report (Bakooshli et al. 2020) this report will launch more detailed studies of muscle function in culture with the potential to develop new therapies for DMD and other muscle disorders.

To further develop the 3D muscle culture, we sought to determine how closely the RMP of 3D muscle resembled native muscle. We report that the mean RMP of healthy muscle is −66 mV at day 8 and −74 mV at day 10 post differentiation This mean RMP for healthy muscle is comparable to a study of membrane characteristics of intact biopsies of intercostal muscle fibers, in which RMP was −76 mV (Gruener et. al., 1979). Ludin (1969) similarly reported a resting potential of −79 mV from excised intercostal muscles and a review shows for most reports of healthy muscle, the resting potential is reported between −73 and −88 mV (Norris, 1976). Therefore, our data from 3D muscle culture align with previously published work of acute measurements from healthy human muscle.

Gruener et. al (1979) also studied human diseased intercostal muscles from biopsies and reported that DMD muscles have a RMP of −57 mV. We report resting potentials from two lines of DMD muscle cultures of −41 and −44 mV. Our result indicates that DMD cells are depolarized compared to healthy cells under identical culture conditions and in agreement with previous reports. We do not yet know why the magnitude of the effect is greater in culture compared to acute biopsy measurements.

Prior work on traditional, two-dimensional, culture systems have also demonstrated that DMD cells grown in culture tend to have a more depolarized resting potential than cells obtained from healthy patients. However, in those reports the resting potential of the healthy cells did not approach that reported for biopsied tissue. For example, Rothman and Bichoff (1983) reported no difference in RMP between healthy and DMD myotubes grown in culture, but the RMP values were −48 and −47.0 mV for the control and Duchenne culture respectively. Further, Merickel et al (1981) report the resting potential for healthy myotubes grown in traditional culture was −53 mV. Altogether, our data indicate that healthy muscle cell tissue grown in 3D culture has an RMP more like that recorded from acute muscle biopsy than traditional culture systems and that DMD tissues have an RMP that is depolarized compared to tissues grown from non-DMD cells.

We also investigated the muscle membrane resistance of 3D cultured muscle and report that DMD muscles possess a lower muscle membrane resistance compared to healthy muscle. Along with the hallmark of DMD showing the detection of creatine kinase leaking from muscle fibers (Ozawa et al., 1999), it has also been shown with electron microscopy studies that there are disruptions being that of lesions on the muscle membrane of DMD fibers (Mokri and Engel, 1975; Pestronk et al., 1982). Greuner et al (1979) reported that membrane resistance measured from DMD muscle biopsy was 1/3 that measured from healthy tissue, and we found a similarly decrease in DMD tissue grown in 3D culture compared to healthy tissue. This would correlate to our finding that the muscle membrane resistance is significantly lower than in healthy muscle.

More recently, studies have been directed at the use of block co-polymers as agents to help with membrane stabilization in *mdx* mice *in vivo*, and *in vitro* (Markham et al., 2015, Houang et al., 2015, Houang et al., 2018). Using block copolymers such as poloxamer188 (P188) have also been shown to protect dystrophic muscles from mechanical stress by stabilizing the phospholipid plasma membrane (Houang et al., 2015). Though various studies have established that P188, an amphiphilic block copolymer, can stabilize the plasma membrane and increase membrane integrity by resealing disruptions of the plasma membrane, these studies have only been conducted in *mdx* mice and human embryonic kidney cells (Andrews and Corrotte, 2018; Bansal et al., 2003; Moloughney and Weisleder, 2012; Kwiatkowski et al., 2020). Since it is known that the membrane of DMD muscles is damaged, we sought out to test the ability of P188 to improve membrane fragility of DMD muscle cells. Incubating DMD muscle tissues with 1% of P188 for 1 hour at 37°C, we found that the RMP of DMD muscles became more negative and we conclude that treatment of P188 helped to improve the electrical properties of the DMD muscle membrane. This suggests that other block co-polymers of different sizes could be looked at and identified as a possible strategy for finding the right copolymer that could get through vessels and patch the fibres further. Using the culture system as a screen to study the effectiveness of membrane repair using different poloxamers could be made possible. Furthermore, the 3D system could also be used for identifying other possible therapeutic reagents that aid in membrane disruption or membrane repair

We were unable to observe active responses using anode break excitation of DMD muscles. Active responses did occur using bridge circuit of some dystrophic muscles, however they were very infrequent and difficult to achieve when compared to the excitability activity of healthy muscles. The difficulty in achieving excitability in the DMD muscles and the decrease in amplitude of the action potentials seen in dystrophic fibers could be due to a decrease in the driving force of Na^+^as well as an increase in Na^+^ channel inactivation in response to a depolarized RMP (Miles et. al., 2011). It was found that human dystrophic intercostal muscles had a 13% reduction in the action potential amplitudes and a 14% reduction in the rise time of action potentials when compared to non-dystrophic muscles (Ludin, 1973). The active responses we found did not follow traditional all or none characteristics tied to reaching thresholds seen in conventional action potentials. Some recordings we would see true action potentials, however this was not frequent and could be due to the maturity of the muscle fibers during culturing and differentiation of our tissue and will require further investigation.

Using transmission electron microscopy to examine the ultrastructure of 3D muscle tissue the key features of skeletal muscle such as striation with different bands were found, however t-tubules with sarcoplasmic reticulum making up the triad were not found. We believe that having a presynaptic component during development of the muscle tissue could allow for further maturation of muscle fibers and could change the presence of such triads. Further maturation of the fibers to consist of these key identifying features of skeletal muscle will be explored in future studies.

In summary, we report the electrophysiological characteristics and ultrastructure 3D bioengineered skeletal muscle tissue *in vitro*. We explore the electrical properties muscle derived from Duchenne muscular dystrophy patients in comparison to healthy muscle. Furthermore, we examine the membrane resistance of DMD muscle cells and test the potential of a co-polymer as a therapeutic agent. Our characterization will form a baseline for further studies directed at skeletal muscle function and serve to establish 3D muscle culture as a pre-clinical platform to test the efficacy of promising candidate therapies for muscle disease

## Acknowledgements

We thank Mohsen Afshar Bakooshli for providing training in the 3D culture system, as well as Doug Holmyard and Ali Dariband at the Nanoscale Biomedical Imaging Facility at The Hospital for Sick Children, Toronto for assistance with transmission electron microscopy. The work was funded by CIHR Project Grant #PJT-168932 to BAS and PMG, and to PMG from the Ontario Institute for Regenerative Medicine and Medicine by Design - a Canada First Research Excellence Program. PMG is the Canada Research Chair in Endogenous Repair.

## Notes

### Competing Interest Statement

The authors have declared no competing interest.

